# Immune gene expression in the mosquito vector *Culex quinquefasciatus* during an avian malaria infection

**DOI:** 10.1101/2022.06.14.496119

**Authors:** L García-Longoria, D Ahrén, A Berthomieu, V Kalbskopf, A Rivero, O Hellgren

**Affiliations:** Department of Anatomy, Cellular Biology and Zoology, University of Extremadura, E-06071 Badajoz (Spain); MIVEGEC (CNRS, Université de Montpellier, IRD), Montpellier, France; Molecular Ecology and Evolution Lab, Department of Biology, Lund University, Sölvegatan 37, SE-22362, Sweden

**Keywords:** Immunogenicity, immunosenescence, *Plasmodium falciparum*, *Plasmodium berghei*, *Anopheles gambiae*, *Anopheles stephensi*

## Abstract

*Plasmodium relictum* is the most widespread avian malaria parasite in the world. It is listed as one of the 100 most dangerous invasive species, having been responsible for the extinction of several endemic bird species, and the near-demise of several others. Here we present the first transcriptomic study examining the effect of *P. relictum* on the immune system of its vector (the mosquito *Culex quinquefasciatus*) at different times post-infection. We show that over 50% of immune genes identified as being part of the Toll pathway and 30-40% of the immune genes identified within the Imd pathway are overexpressed during the critical period spanning the parasite’s oocyst and sporozoite formation (8-12 days), revealing the crucial role played by both these pathways in this natural mosquito-*Plasmodium* combination. Comparison of infected mosquitoes with their uninfected counterparts also revealed some unexpected RNA expression patterns earlier and later in the infection: Significant differences in expression of several immune effectors were observed as early as 30 minutes after the ingestion of the infected blood meal. In addition, in the later stages of the infection (towards the end of the mosquito lifespan), we observed an unexpected increase in immune investment in uninfected, but not in infected, mosquitoes. In conclusion, our work extends the comparative transcriptomic analyses of malaria-infected mosquitoes beyond human and rodent parasites and provides insights into the degree of conservation of immune pathways and into the selective pressures exerted by *Plasmodium* parasites on their vectors.

## INTRODUCTION

Malaria parasites are one of the most successful parasites on the planet. They are largely known for infecting humans, where they are responsible for an estimated 600 000 deaths per year (Health and Who 2021). These protozoans can also be found infecting hundreds of other terrestrial vertebrate species, including non-human primates, ungulates, rodents, bats, birds, lizards and snakes. There are currently almost a thousand different *Plasmodium* species, with more being described every year. But for a few minor differences, all these species share a nearly identical life cycle, with an asexual replicative stage in the vertebrate host, and a sexual stage in a dipteran vector, usually a mosquito (Eckhoff 2011; Valkiunas and Iezhova 2018).

The importance of mosquitoes in malaria transmission has made them particularly important targets of research (Kar et al. 2014; Ryan et al. 2020; Keleta et al. 2021). Research into the development and testing of new insecticides is ongoing but the evolution of insecticide resistance has emerged as a serious challenge to malaria control efforts (Moyes et al. 2021). As a result, a great deal of effort is currently focused on alternative strategies aimed at curbing malaria transmission. One such promising approach is the potential to harness mosquito physiology and immunity for the development of transmission-blocking interventions (Wadi et al. 2018; Challenger et al. 2021). *Plasmodium* parasites go through multiple stages inside mosquitoes. Mosquitoes ingest a bloodmeal containing female and male *Plasmodium* gametocytes, which fuse to become a motile ookinete that traverses the midgut wall to form an oocyst. Oocysts undergo repeated rounds of mitosis to create a syncytial cell with thousands of nuclei. In a massive cytokinesis event, thousands of haploid daughter sporozoites are liberated into the haemolymph, where the infective sporozoites migrate to the mosquito salivary glands for transmission to a new host host (Valkiūnas 2005; Howick et al. 2019).

The mosquito immune system has received a lot of attention as mounting evidence points to its critical role in eliminating a sizeable proportion of parasites invading the midgut epithelium (Hajkazemian et al. 2021). In recent years, several genomic and transcriptomic studies performed on the main vectors of human malaria, the *An. gambiae* species complex, have allowed the identification and quantification of numerous transcripts involved in the mosquito immune response to a malaria infection (Reynolds et al. 2020; Carr et al. 2021). There are two main arms of the mosquito innate immune response against *Plasmodium*: the humoral response involving the transcriptional regulation of anti-microbial peptides (AMPs), and the cellular response, that includes phagocytosis and/or melanization (Clayton et al 2014). Three main signalling pathways have been shown to be involved in the immune cascade that ultimately leads to the destruction of the parasite: the Toll, the immune deficiency (Imd) and the Janus kinase signal transducer of activation (JAK-STAT) pathways (Tikhe and Dimopoulos 2021). These pathways involve different complex immune cascades that ultimately allow the REL1 (Toll), REL2 (Imd) or STAT (JAK-STAT) transcription factors to enter the nucleus and transcriptionally activate immune effector genes, such as AMPs, complement factors and Nitric Oxide Synthase (Clayton et al 2014). These transcription factors are negatively regulated in the cytoplasm by Cactus (Toll), Caspar and Caudal (Imd) and SOCS and PIAS (JAK-STAT, Clayton et al 2014).

One key insight from transcriptomic studies has been that natural and artificial mosquito-*Plasmodium* combinations (i.e. combinations that are found in the wild, or not) have significantly different immune activation profiles (Sreenivasamurthy et al 2013). For example, rodent *P. berghei* parasites have a greater impact on the transcriptome of the human malaria vector *A. gambiae*, regulating hundreds of genes belonging to different functional classes, than human *Plasmodium falciparum* parasites (Dong *et al* 2006). Most notably, in *A. gambiae* the most effective pathway for the defense against *P. berghei* is the Toll pathway, while for the defense against *P. falciparum* it is the Imd pathway (Garver et al. 2009; Clayton et al. 2014). This suggests that the rodent malaria model may be less relevant to the study of the immune strategies employed by malaria-infected mosquitoes than a *Plasmodium*-mosquito combination with a long co-evolutionary history.

Avian malaria has played a key role in the development of knowledge of human malaria parasites, including the elucidation of key aspects of the biology and transmission of *Plasmodium*, and the routine testing and development of the first vaccines and antimalarial drugs (Rivero and Gandon 2018). Here we present the first transcriptomic analysis of mosquito immune genes in response to a natural avian malaria infection. For this purpose, we use *Plasmodium relictum*, one of the most widespread avian malaria parasites in the world, (Kazlauskiene et al. 2013, Valkiūnas et al. 2018). This species is responsible for several bird species extinctions (Atkinson and Samuel 2010), threatening several others (Butchart 2008), and is currently listed as one of the 100 most dangerous invasive species (Lowe et al 2000). Its main natural vectors are mosquitoes of the *Culex pipiens* complex, comprising *Culex pipiens* and *Culex quinquefasciatus* (Fonseca et al 2004). The genome of *Culex quinquefasciatus*, which also vectors human filarial parasites and the West Nile virus, was sequenced a decade ago (Arensburger et al., 2010). Since then, the number of studies investigating the transcriptomic response of this species to a pathogen infection has been surprisingly limited (Bartholomay et al. 2010; Girard et al. 2010; Shin et al. 2014).

We investigate the temporal gene expression patterns of *Cx quinquefasciatus* mosquitoes infected with *P. relictum* focusing on immune genes that have been previously identified as being key for the immune response in human vectors of malaria. For this purpose, we compare the transcriptome of infected and control mosquitoes at four different phases of the infection, each corresponding to a key stage of the parasite’s development: within the first 30 min of the infected blood meal (gametocyte activation and formation of gametes), and 8 days (peak oocyst production), 12 days (peak sporozoite production) and 22 days (ending stages of the infection) later. The full transcriptomic analyses of the parasites at each of these different stages has been published in a separate paper (Sekar et al. 2021). We compare our results to previous work carried out in other mosquito-*Plasmodium* and *Culex*-pathogen combinations. Our study extends the comparative transcriptomic analyses of malaria-infected mosquitoes beyond human and rodent parasites and provides insights into the degree of conservation of immune pathways and into the selective pressures exerted by *Plasmodium* parasites on their vectors.

## RESULTS

### Principal component analysis

A principal component analysis (PCA) was run for each group of genes (Toll, Imd and JAK/STAT pathways, Figure 1). The first (PC1) and second (PC2) axes explain 59% and 13% of variance in the expression of genes of the Toll pathway (Figure 1A), 62% and 18% of the Imd pathway (Figure 1B) and, 66% and 21% of the JAK/STAT pathway (Figure 1C). The PCA analyses show differences in the global expression patterns of infected and non-infected mosquitoes across the four different time points in the three immune pathways (Figure 1). The largest differences in scores across both the PC1 and PC2 are seen in control mosquitoes at 30mpi. A certain discrimination based on scores of PC was also observed in control mosquitoes at 8 and 12 dpi. Interestingly, there was no clear discrimination between the expression levels of infected mosquitoes across different sampling times, except for genes in the Toll pathway of 30mpi mosquitoes which cluster separately from their control counterparts (Figure 1A).

**Figure 1.**
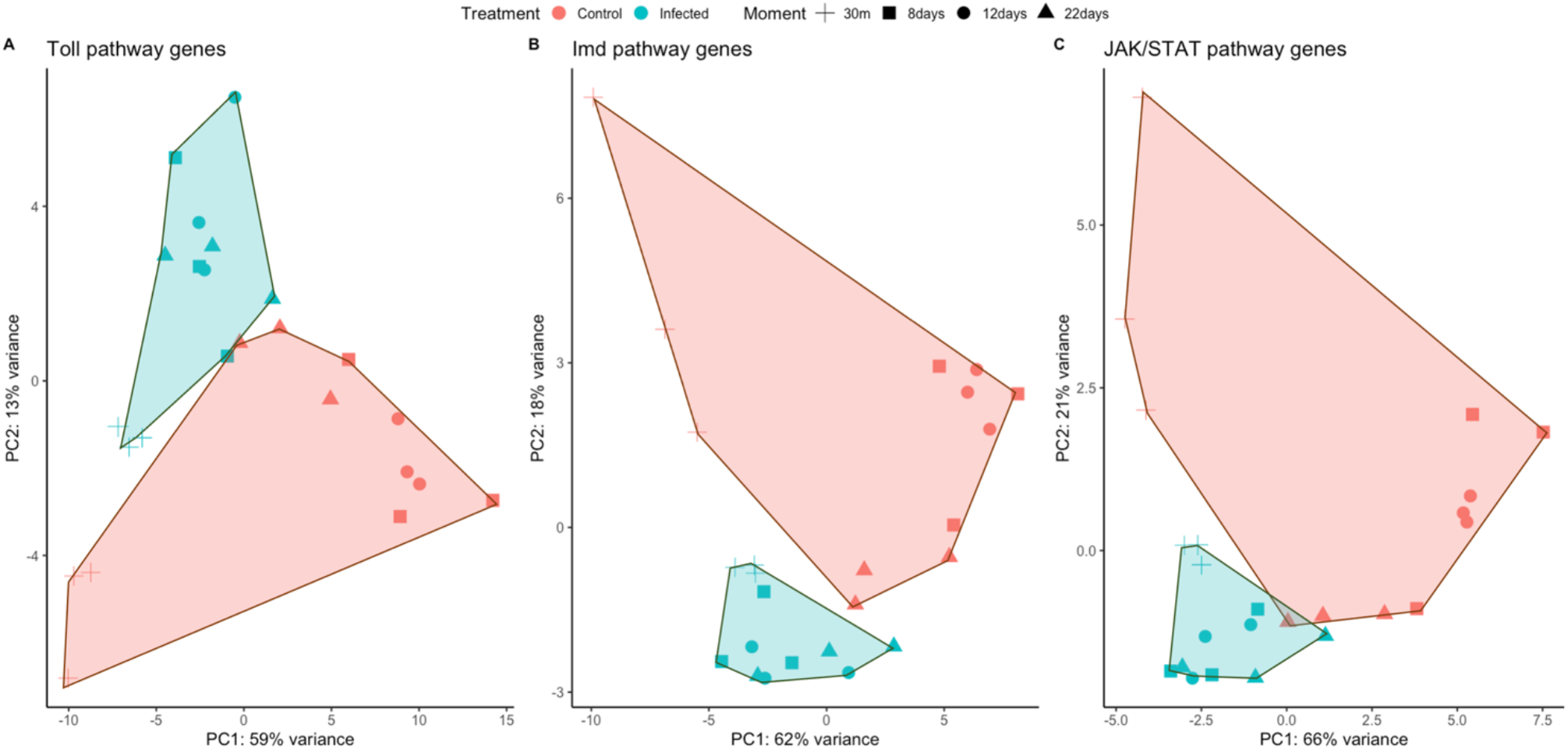
Principal component analyses of expression levels from the three immune pathways (A: Toll, B: Imd, C: JAK/STAT). Results have been colour marked depending on the mosquito condition: red infected; blue non infected. Also, different sampling times are shown with different shapes.

### Overall immune gene expression pattern

Our analyses detected 92 immune genes belonging to the Toll pathway, 74 to the Imd pathway and 42 to the JAK/STAT pathway. The number of differentially expressed genes differed widely between pathways and sampling points (Figure 2 – see below). In particular, there is a striking up regulation of Toll pathway genes at 8 dpi and 12 dpi, and a near absence of DEGs towards the end of the infection (22 dpi).

**Figure 2.**
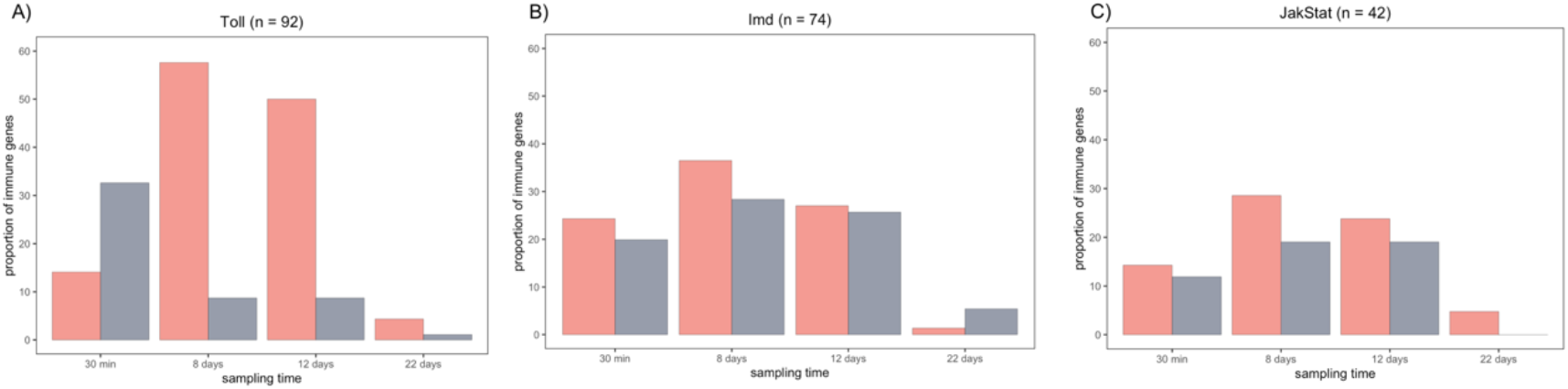
Proportion of differentially expressed genes (DEG) that are upregulated (pink) or downregulated (grey) at each sampling point after infection with *P. relictum*. The name of the pathway and total number of genes identified for each pathway are given in the header.

We classed the DEGs into four different functional classes: receptors, transcription factors, inhibitors and effectors, and the magnitude of gene expression differences between infected and uninfected mosquitoes was represented as log-fold change (LFC) values (Figure 2). The highest overall LFC values are observed within the Toll pathway. Many of the receptors, transcription factors and effectors within the Toll and Imd pathways followed a similar pattern: they were higher in infected mosquitos at 8 and 12 days dpi than earlier (30 mpi) and later (22 dpi) in the infection (Figure 2 but also see Supplementary Table 1). This pattern was, however, less apparent within the JAK/STAT pathway (Figure 2C). Four of the 7 inhibitor genes identified were under-expressed while two others (one Imd and one JAK/STAT) were significantly over-expressed at 8 dpi and 12 dpi (Figure 2).

### Biological function of differentially expressed genes

We identified a significant up-regulation of genes related to Imd and Toll pathways (Figure 3A, Supplementary Table 2). Most of the present genes in the Toll and Imd pathways showed either significant up or down regulation during the different infection phases. For JAK/STAT pathway most of the genes detected were down regulated or not DEG (Figure 3B, Supplementary Table 2). The biological function of the genes within the Imd, Toll and JAKSTAT pathways are shown in Figure 4. The pattern of up or down regulation of these different genes are discussed below separately for each sampling time.

**Figure 3.**
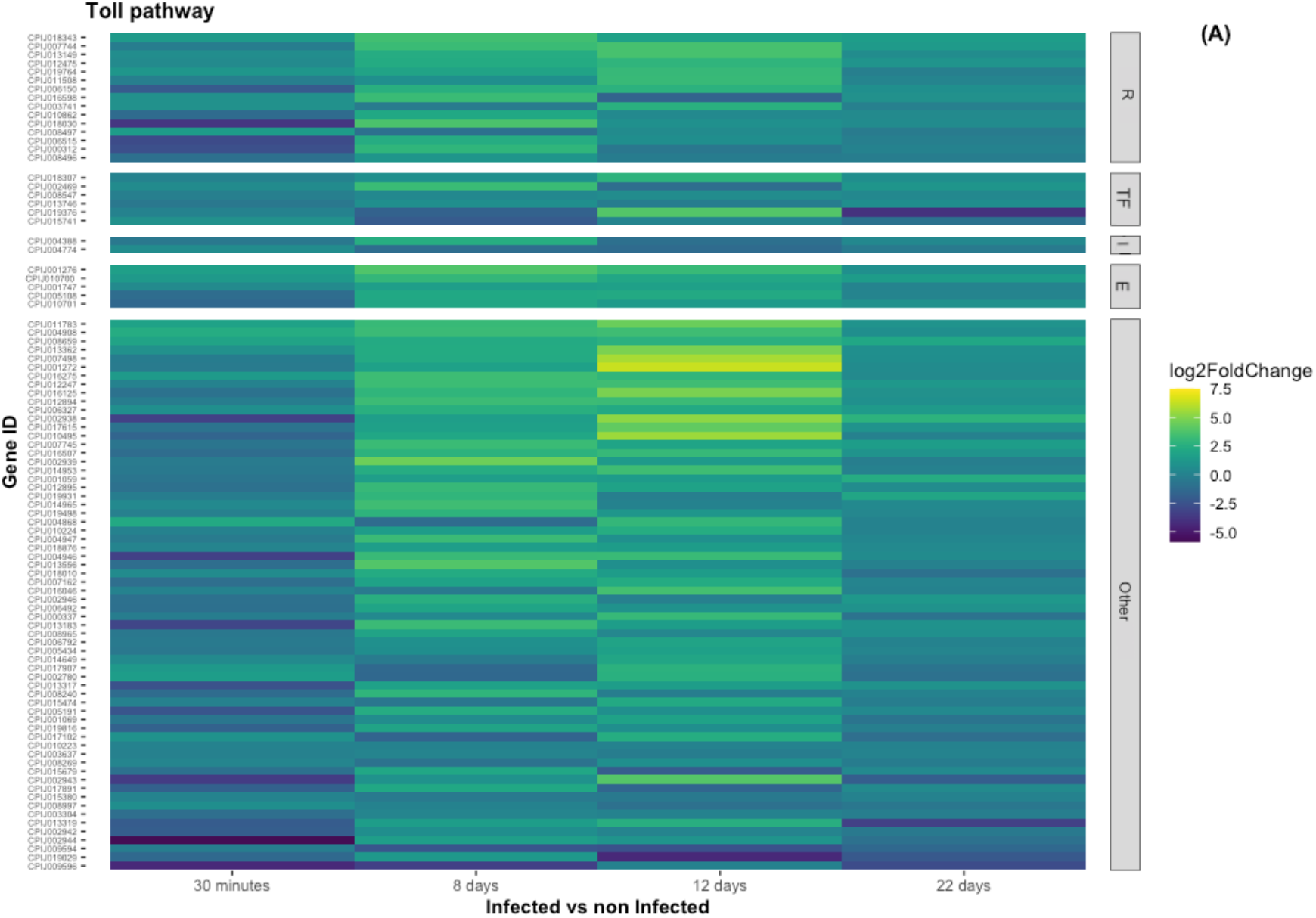
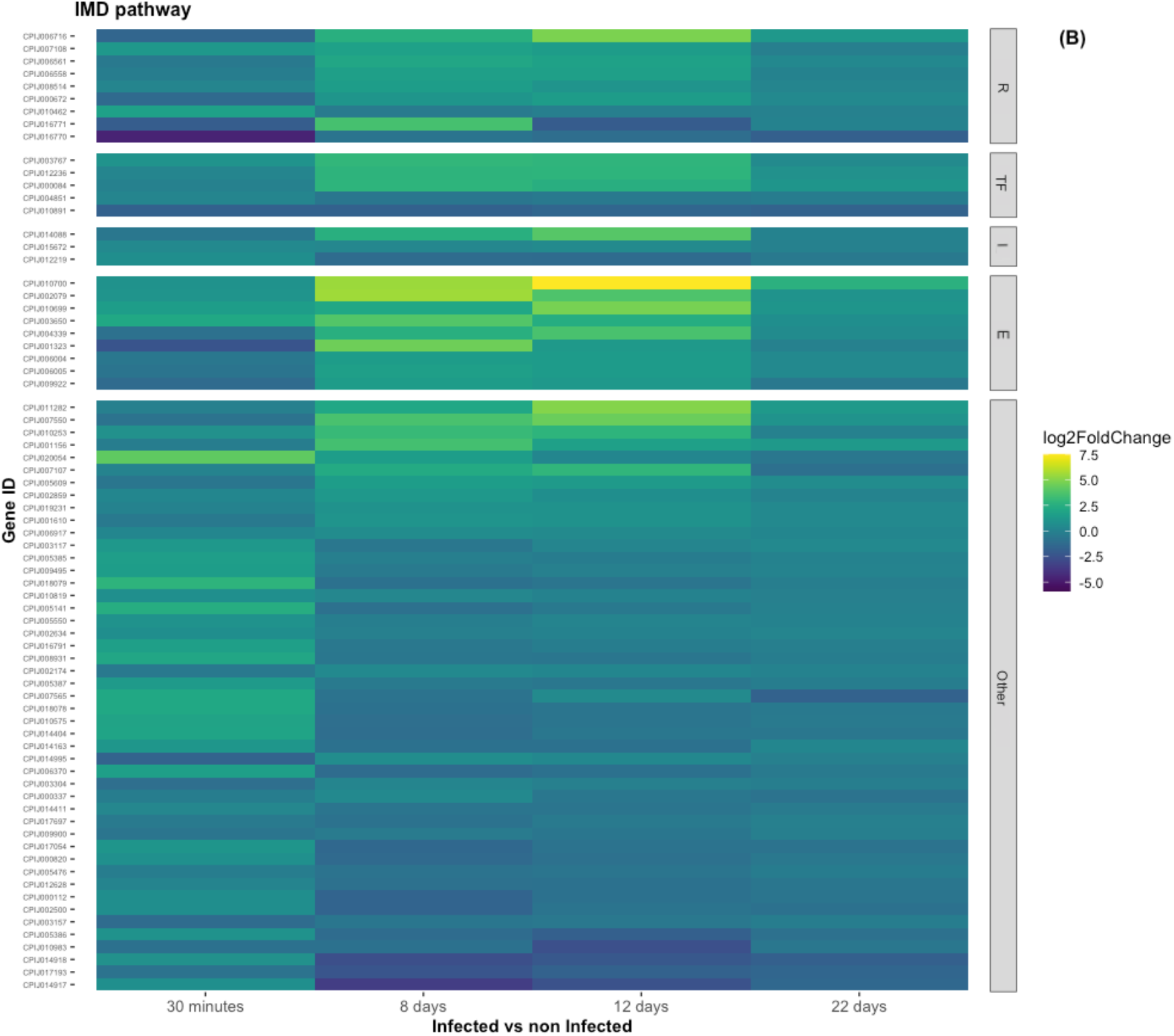
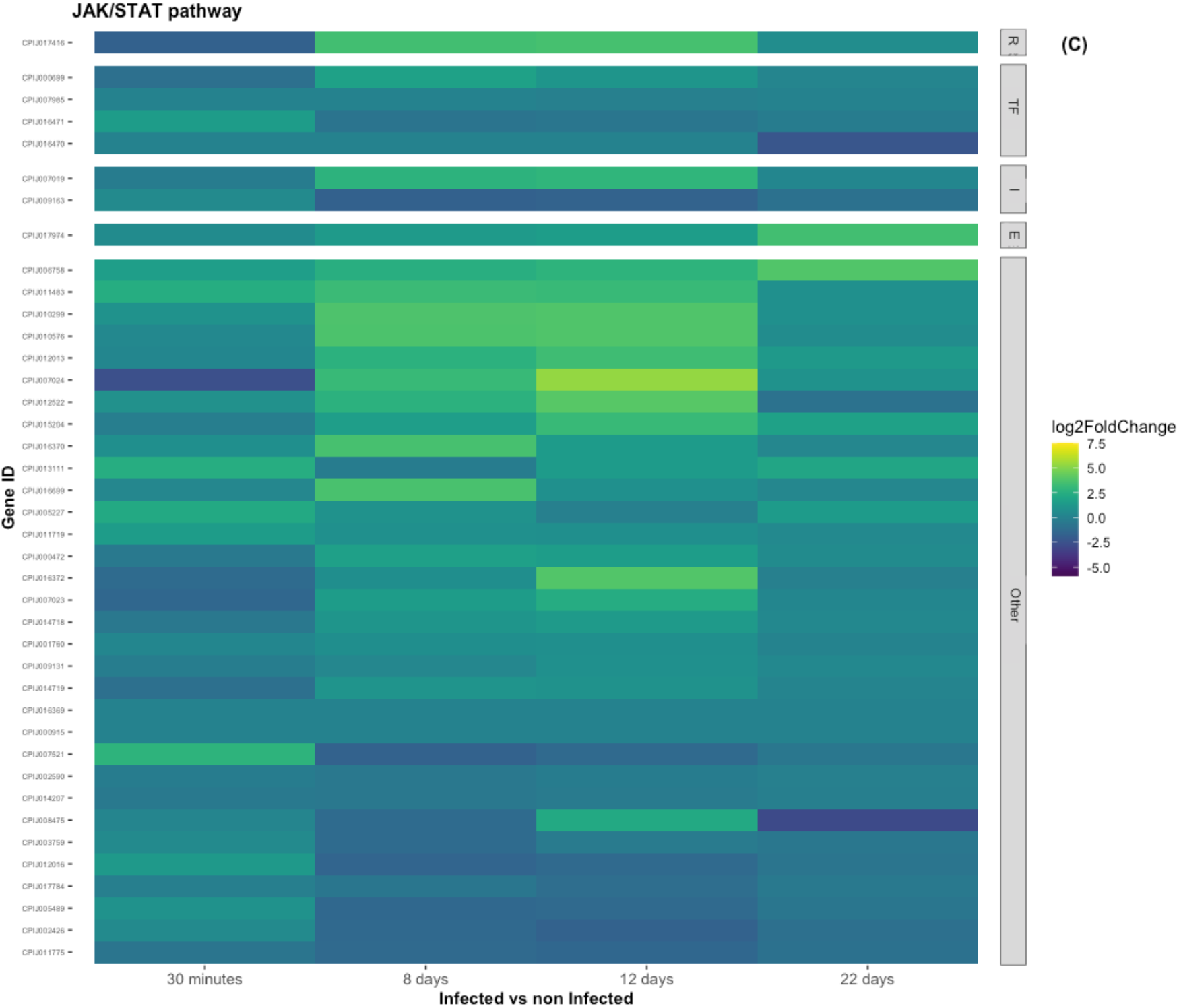
Differentially expressed genes (DEGs) at 8 dpi (peak oocyst) and 12 dpi (peak sporozoite) mapped onto the KEGG Toll and Imd signalling pathways of *Drosophila melanogaster* (Kanehisa et al. 2021) (A), and JAK/STAT pathway adapted from the *Homo sapiens sapiens* pathway (Kanehisa et al. 2021) (B). Green: genes upregulated in infected mosquitoes as compared to control ones. Red: genes down regulated. Grey: no significant difference in expression but with observed expression in the data. White: genes not present in our analysis. Yellow: genes detected but with 0 transcripts (TPM) in all sampling times. Genes with N/A values are not plotted (See supplementary Table 1 but also Table 2)

**Figure 4.**
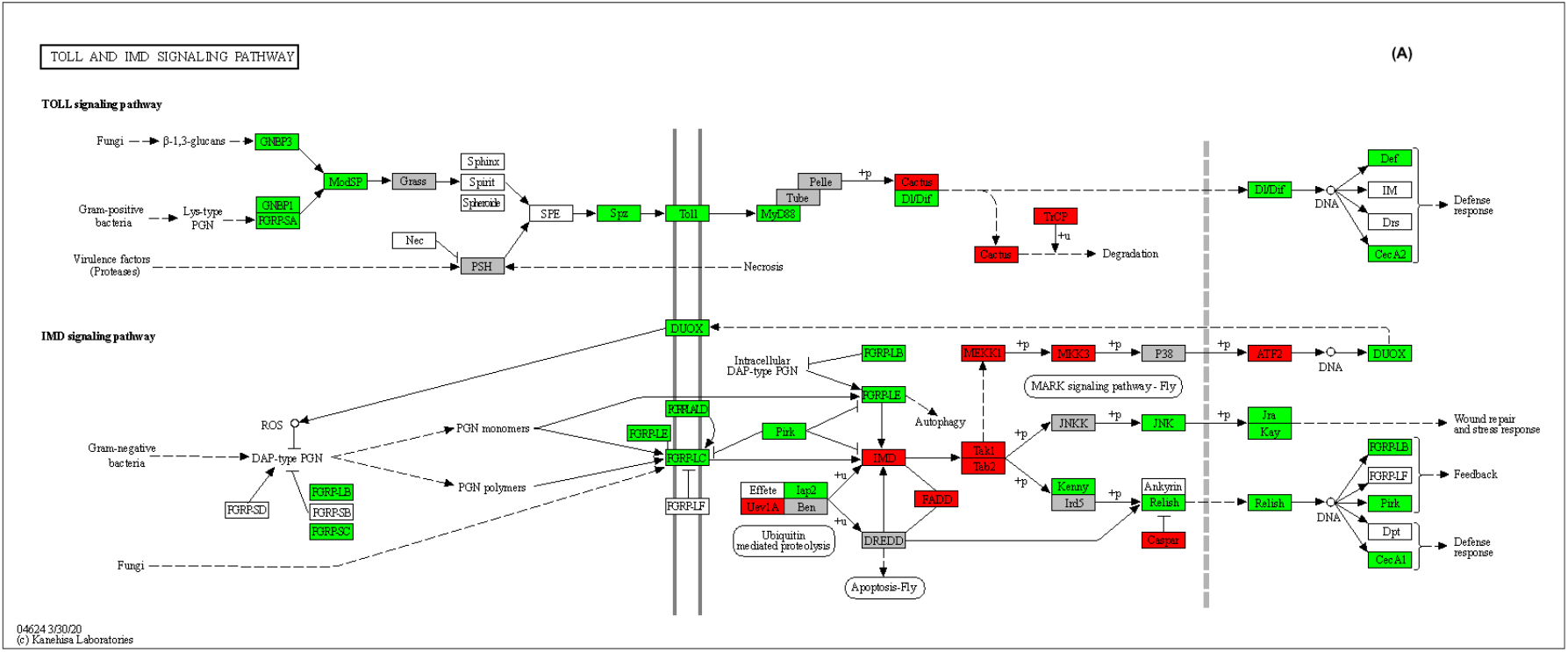
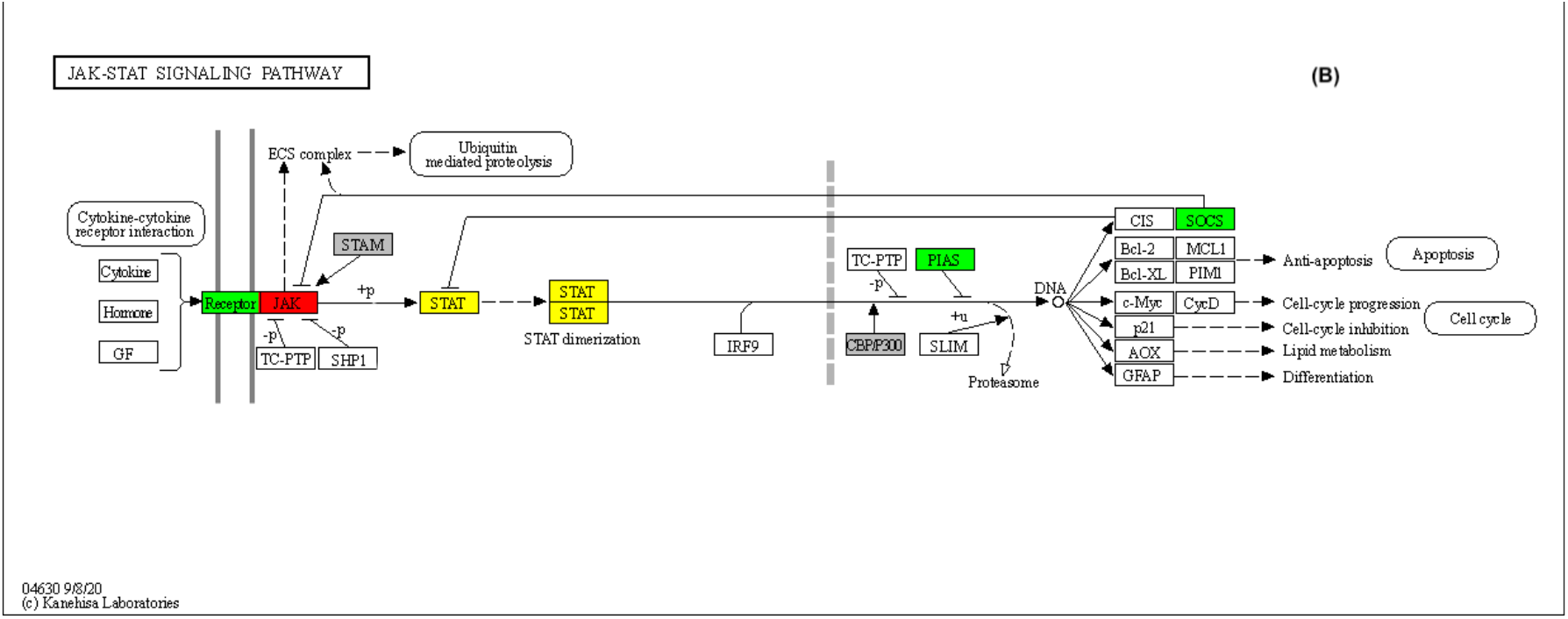
Heatmaps for the different pathways showing Log-fold change (LFC) values resulting from the statistical comparison of infected versus control mosquitos at each sampling time (A: Toll, B: Imd, C: JAK/STAT). LFC have been classified as follows: Receptors (‘R’), Transcription factors (‘TF’), Inhibitors (‘I’), Effectors (‘E’) and ‘Other’ (grouping all other genes within the pathway).

#### 30 minutes post-blood meal

Only two Toll receptors (CPIJ019764, CPIJ018343, Figure 5A) were significantly upregulated 30 minutes after the ingestion of an infected blood meal.

**Figure 5.**
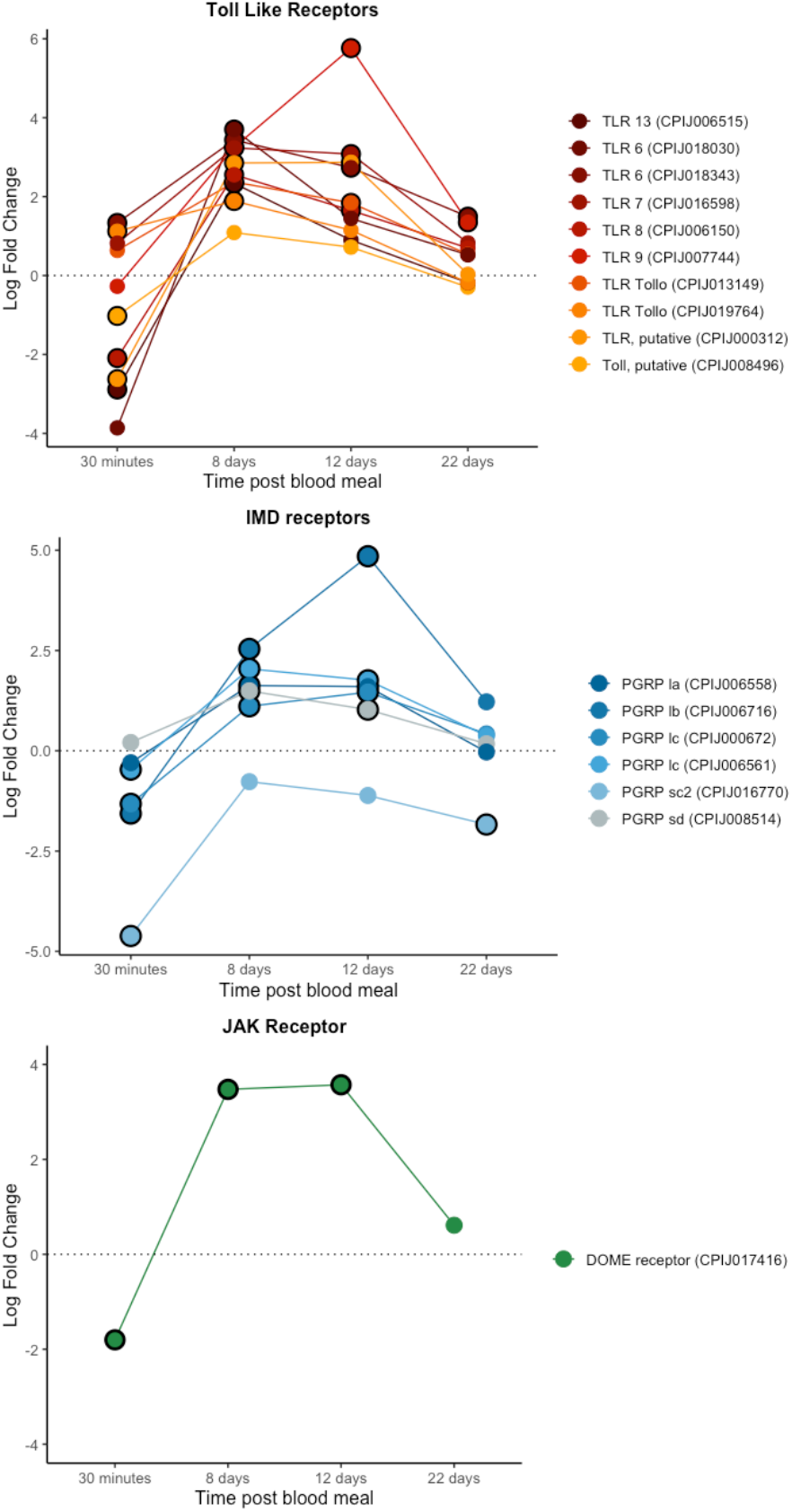
Differential expression pattern of receptor genes between infected and control mosquitoes across different sampling points. Differential expression is represented as log-fold change (LFC) values. A: Toll pathway, B : Imd pathway, C: JackStat pathway. Values above the dashed horizontal line indicate a higher expression in infected than uninfected mosquitoes. The black circle around each dot indicates a significant difference when comparing infected and control mosquitos (adjusted p < 0.05).

We did not detect any up regulated transcription factors in either the Toll, Imd or JAK/STAT pathways (Figure 6). In contrast, there was a down-regulation of the Imd transcription factor (CPIJ010891) in infected mosquitos (Figure 6B). No inhibitors were significantly up or down regulated for any of the 3 pathways at this time point. Interestingly, some effectors as defensin-A and Cecropin-A were up-regulated as early as 30 minutes post infection, while others as cecropin genes (A2 and B1) were significantly down-regulated (Figure 7A). We did not detect any significant difference in the expression of NOS (Figure 7B).

**Figure 6.**
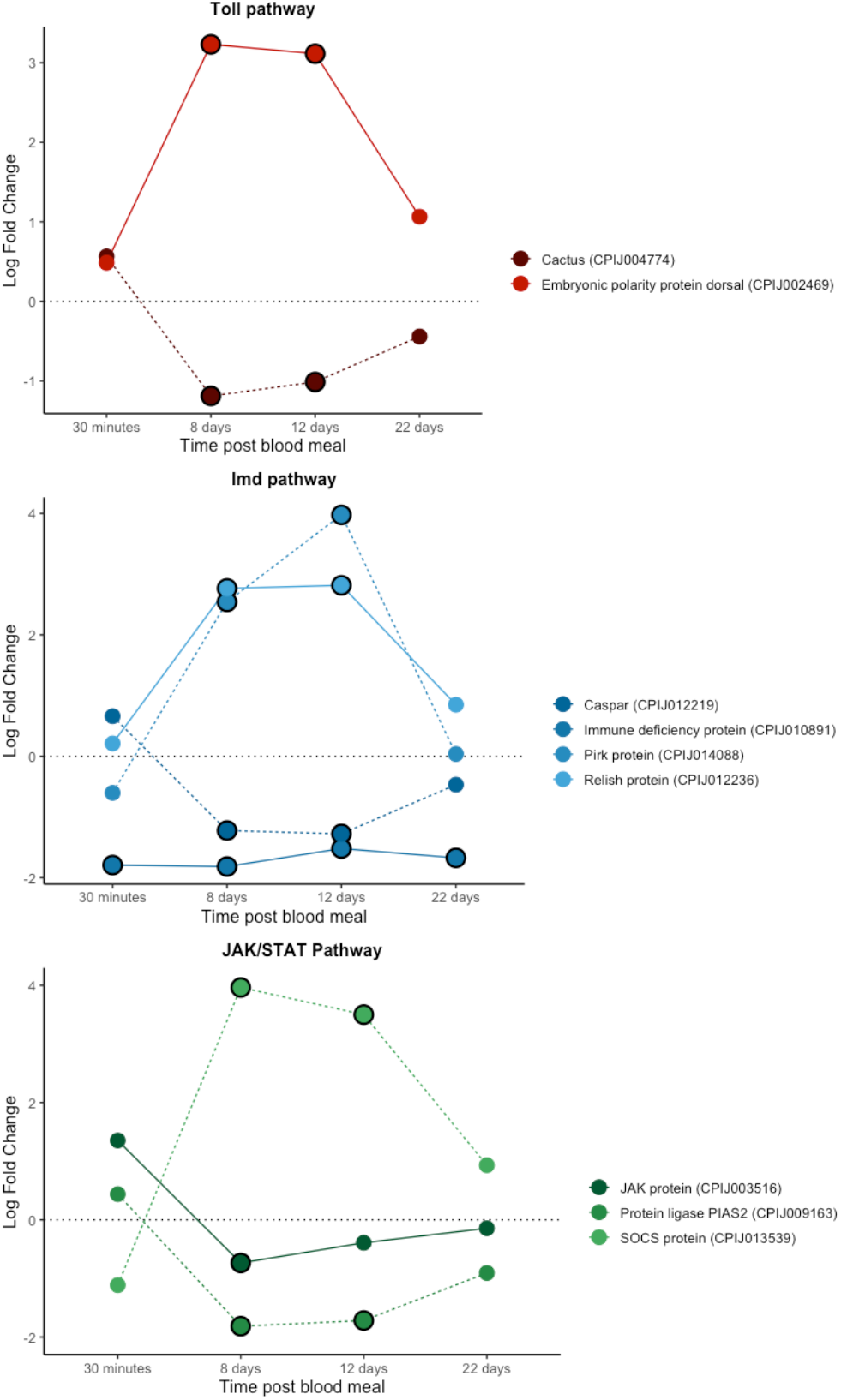
Differential expression pattern of transcription factor and inhibitor genes between infected and control mosquitoes across different sampling points. Differential expression is represented as log-fold change (LFC) values. A: Toll pathway, B : Imd pathway, C: JAK/STAT pathway. Each colour represents a transcription factor or inhibitor combination. Dark tones and continuous lines indicate transcription factors and soft tones and dashed lines indicate inhibitor genes. Note that each transcription factor has the same colour that its inhibitor (blue, orange or purple). The black circle around each dot indicates a significant difference when comparing infected and control mosquitoes (adjusted p < 0.05).

**Figure 7.**
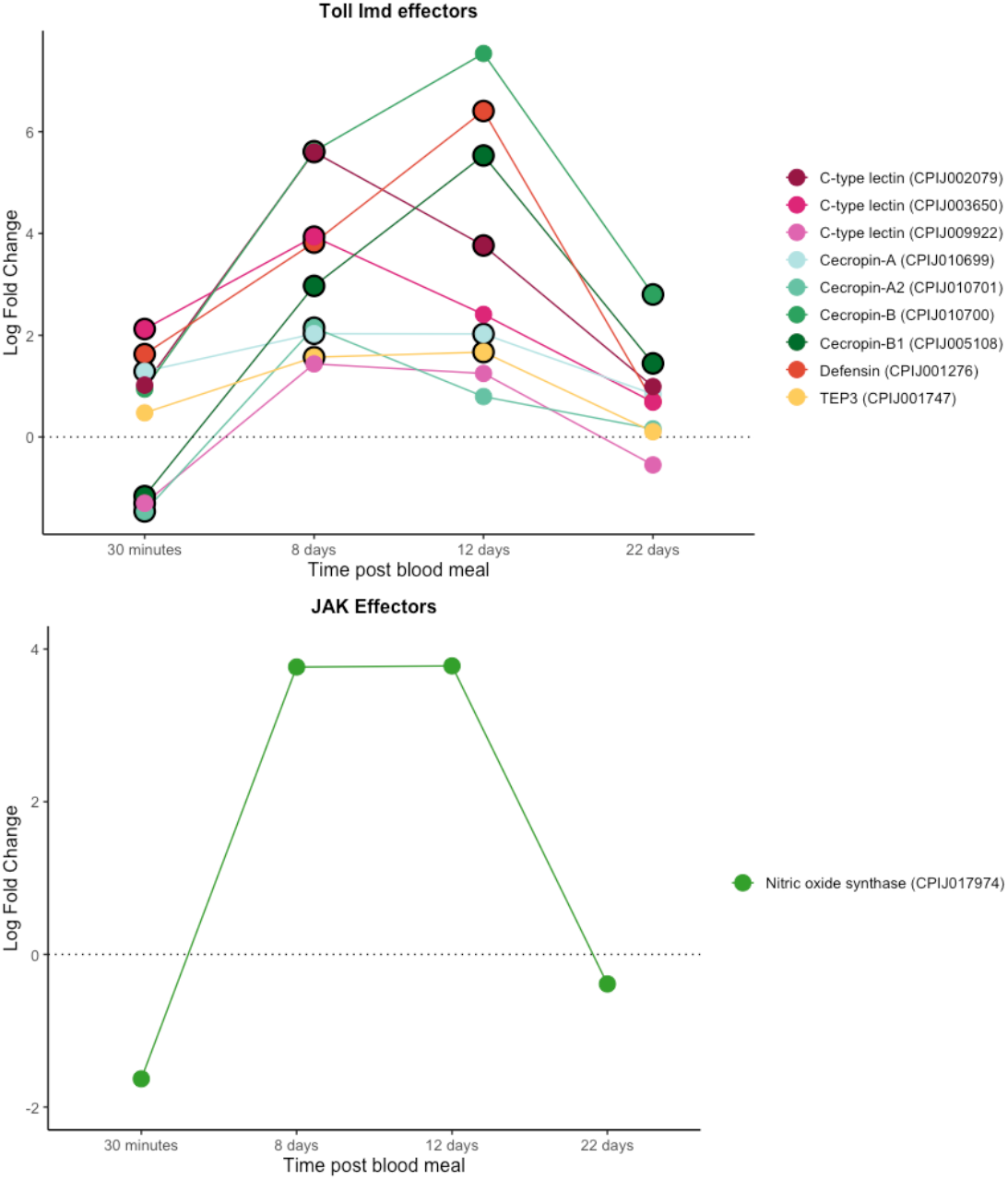
Differential expression pattern of effector genes between infected and control mosquitoes across different sampling times. Differential expression is represented as log-fold change (LFC) values. A: effectors for Toll and Imd pathways; B: NOS. The black circle around each dot indicates a significant difference when comparing infected and control mosquitos (adjusted p < 0.05).

Genes of PPO cascade were not significantly up regulated at this sampling time (Figure 8).

**Figure 8.**
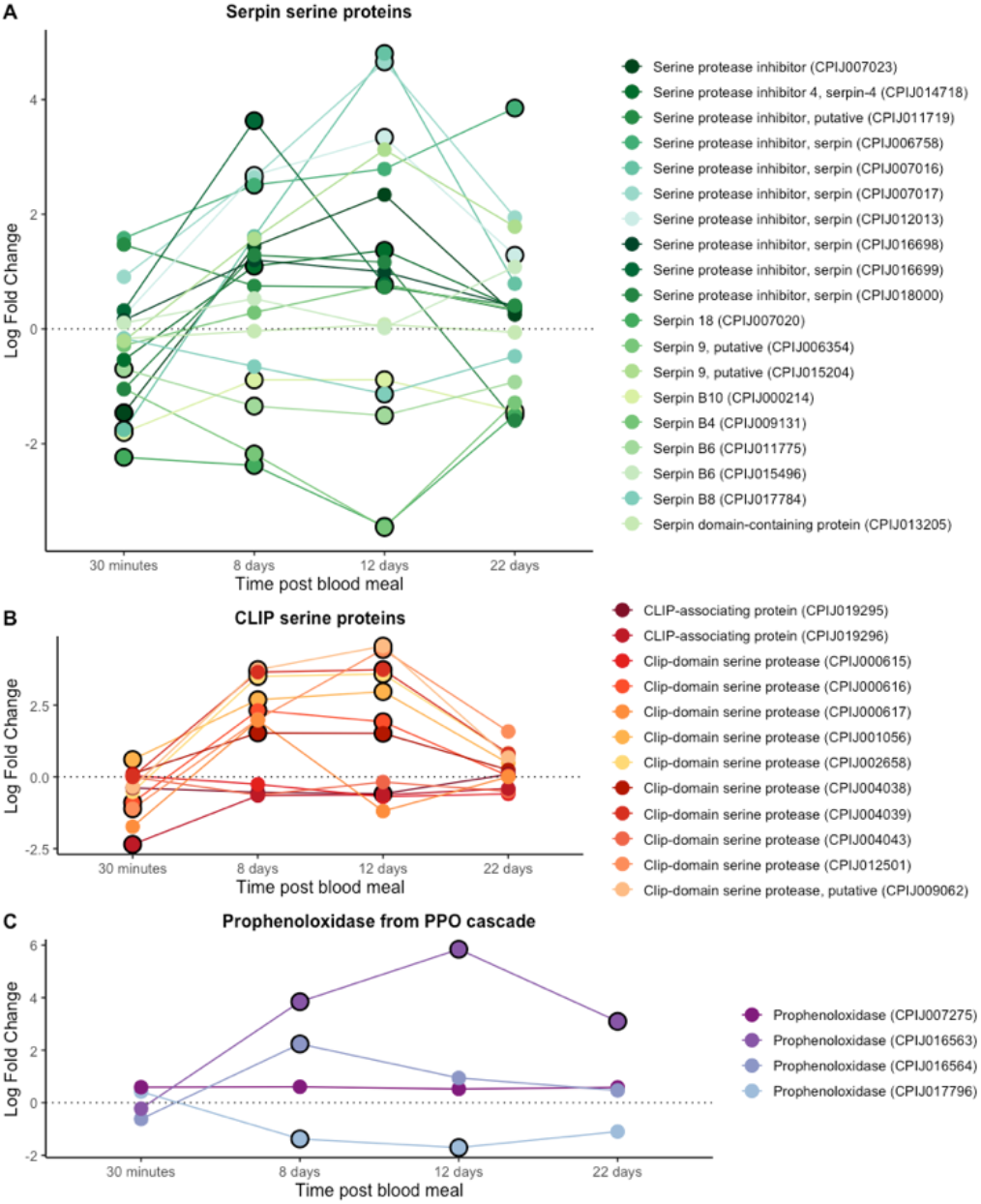
Differential expression pattern of (A) Serpin serine proteins; (B) CLIP serine proteins and (C) Prophenoloxidase fom PPO cascade. Differential expression is represented as log-fold change (LFC) values. The black circle around each dot indicates a significant difference when comparing infected and control mosquitos (adjusted p < 0.05).

#### 8 days and 12 days post blood meal

Day 8 post blood meal is associated to a significant upregulation of receptors in the Toll, Imd and JAK/STAT pathways (Figure 5). The level of expression of these receptors did not change significantly between 8dpi and 12dpi, except for TLR9 (CPIJ007744) in the Toll pathway and PRPlb (CPIJ006716), which showed a further increase.

The Dorsal transcription factor within the Toll pathway (CPIJ002469) was differentially up-regulated at 8 and 12 days while its inhibitor, cactus protein (CPIJ004774), was significantly down regulated (Figure 6A). A similar pattern was observed with the relish transcription factor (CPIJ012236) and its caspar inhibitor (CPIJ012219) within the Imd pathway (Figure 6B). In contrast, an opposite pattern (downregulation of the transcription factor, upregulation of the corresponding inhibitor) was observed for the Immune deficiency protein transcription factor (CPIJ010891) and Pirk inhibitor (CPIJ014088) in the Imd pathway (Figure 6B), and the Signal transducing adapter molecule 1 (CPIJ003516) and its SOCS inhibitor (CPIJ013539, Figure 6C).

As expected, most antimicrobial peptides and effectors were significantly upregulated at 8 and 12 dpi (Figure 7A). NOS showed a similar pattern, showing a drastic increase between 30mpi and 8dpi, even though the differential expression levels did not reach statistical significance at any of the time points (Figure 7B).

CLIP serine proteases and prophenoloxidases from the PPO cascade were significantly up regulated at this sampling time (Figure 8) while some serpin proteins remained significantly down regulated (Figure 8C).

#### 22 days post blood meal

Receptor expression decreased drastically between the 12 and 22 dpi time points. Two Toll receptors (CPIJ018343 and CPIJ007744) were significantly up-regulated 22 days after infection (Figure 5A). One of the receptors in the Imd pathway (CPIJ016770), however, was significantly downregulated.

Neither the Toll transcription factor nor its inhibitor protein were differentially expressed at 22 days post infection (Figure 6A). The same results were observed for Relish and Caspar within the Imd pathway (Figure 6B). The Immune deficiency transcription factor was significantly downregulated in infected mosquitoes throughout the course of the infection (Figure 6B).

Antimicrobial peptide and NOS expression decreased drastically between the 12 and 22dpi time points. Only cecropin-B and B1 (CPIJ0005108, CPIJ010700) were significantly up-regulated throughout the infection (Figure 7A).

Any CLIP serine proteins or prophenoloxidases were significantly up regulated except for one prophenoloxidases (CPIJ007275) (Figure 8). Most serpin genes were not significantly expressed (Figure 8A).

Our results showed that the lack of differential expression between infected and uninfected mosquitoes was due to an increase in expression in uninfected mosquitoes rather than a decrease in expression in infected ones (Figure S1). The overall pattern was that the infected mosquitoes increased their gene expression at day 8 and that gene expression was kept until day 22, whereas the uninfected mosquitoes had a new set of genes being upregulated at day 22 (Figure S1C Figure S1F).

## DISCUSSION

*Cx quinquefasciatus* is an epidemiologically important vector of an exceptionally diverse array of taxonomically different pathogens, including arboviruses (West Nile, St. Louis encephalitis and Rift Valley viruses), filarial worms (*Wuchereria bancrofti*) and protozoans (avian *Plasmodium* and avian *Trypanosoma*). Although *Culex* mosquitoes diverged from their *Anopheles* and *Aedes* counterparts during the early Jurassic (∼ 160-200 million years ago) and early Cretaceous (∼ 130 million years ago) periods, respectively (da Silva et al. 2020; Lorenz et al. 2021) they still share pathogen groups for which they act as vectors for e.g. *Plasmodium* spp. The first *Cx quinquefasciatus* genome sequence revealed a significant expansion in the number of immune genes compared with *Anopheles gambiae* (+120 genes) and *Aedes aegypti* (+83 genes, (Bartholomay et al. 2010)) possibly reflecting the higher diversity of pathogens to which this species is exposed both during its larval and adult lives. While in the last decade the number of transcriptomic studies identifying the immune pathways activated in *Anopheles* and *Aedes* mosquitoes in response to malaria and arboviral infections has grown exponentially (Cirimotich et al. 2010; Ruiz et al. 2019; Vargas et al. 2020; Singh et al. 2021), we still know comparatively little about how *Culex* mosquitoes respond to infections (but see (Girard et al. 2010)).

The transcriptomic analyses of *Cx quinquefasciatus* infected with *P. relictum*, a highly prevalent (Hellgren et al. 2015) and often virulent (Bueno et al. 2010; Cellier-Holzem et al. 2010) avian malaria parasite, reveals dynamic changes in genes in the Toll, Imd and, to a lesser extent, JAK-STAT immune pathways throughout the infection. Previous work on the key human malaria vector, *An. gambiae*, has shown that while the Toll pathway is particularly effective against rodent *P. berghei* parasites, human *P. falciparum* parasites are largely controlled via the Imd pathway (Clayton et al 2014). The reasons for this are not yet clear, although these striking differences in immune response have sounded a cautionary note about the dangers of interpreting the transcriptomes of non-natural mosquito-*Plasmodium* combinations (Boëte 2005).

Here we show that in *Cx quinquefasciatus* infected with *P. relictum*, over 50% of immune genes identified as being part of the Toll pathway and 30-40% of the immune genes identified within the Imd pathway are overexpressed during the critical period spanning oocyst and sporozoite formation (8-12 days), revealing the crucial role played by both these pathways in natural mosquito-*Plasmodium* combinations. A significant upregulation of Toll-like receptors (TLRs) and Peptidoglycan recognition receptors (PGRRs) is observed. In addition, key transcription factors within the Toll (Dorsal) and Imd (Relish) pathways are significantly higher in infected vs control mosquitoes, while their corresponding inhibitors (Cactus and Caspar, respectively) are significantly lower. A wide range genes controlling immune effectors are also significantly more expressed in infected vs control mosquitoes 8-12 days after an infected blood meal, including several genes which codes antimicrobial peptides which previously have been shown to play an important role in the clearance of rodent and human *Plasmodium* (Cecropin A, Cecropin B and Defensin, Kokoza et al. 2010, Raulf et al. 2019, Simões et al. 2022). NOS, the enzyme responsible for the production of nitric oxide, which has been shown to be linked to *Plasmodium* killing in the midgut (Luckhart et al. 1998; Kumar et al. 2003; Peterson et al. 2007) was also overexpressed in infected relative to uninfected mosquitoes although the differences were not statistically significant.

*Plasmodium* oocysts are vulnerable to melanization, a cascade that initiates with the proteolytic activation of the prophenoloxidase (proPO) and which is controlled by a series of CLIP-domain serine proteases and their serpin inhibitors (Zhang et al. 2016). Two CLIP serine proteases, CLIPB8 and CLIPB9, and a one serpin, SRPN2, have been identified as crucial elements of the melanization cascade in *An. gambiae* mosquitoes infected with human malaria. Several other CLIPB proteinases, including CLIPB1, 3, 4, 14, and 17 also affect rodent malaria parasite (Cao et al. 2017). At 8-12 days post-infection we observed an over-expression of several proPOs, and CLIP-domain serine proteases, and an under-expression of serpins, indicating that the phenoloxidase (melanization) cascade may play an important role in the defense against *P. relictum*. At this point, we do not know which, if any, of the CLIPs and serpins triggered in *Cx. pipiens* in response to a *Plasmodium* infection play a role in the mosquito melanization cascade.

An unexpected result was the significant differences in expression of several immune effectors as early as 30 minutes after the ingestion of the infected blood meal. Although the exact time course of the onset of an avian malaria infection in mosquitoes has not yet entirely elucidated (Rivero and Gandon 2018), extrapolations from human and rodent malaria would suggest that at this time point the gametocytes ingested with the blood meal are transforming to gametes before fusing to produce a motile zygote within the blood bolus (Tahar et al. 2002; Dong et al. 2006). Several immune effectors (Cecropine A, Defensin and C-type lectin) are significantly over expressed in infected mosquitoes at this early time point. Several non-exclusive explanations can be proposed for this observed phenomenon. First, gametes or zygotes may be immunogenic. Several proteins (e.g. P25/P28), which are transcriptionally repressed in the gametocyte, are expressed as soon as the gametes are formed (del Carmen Rodriguez et al. 2000). These proteins are highly immunogenic for the vertebrate immune system (Saxena et al. 2007), but whether they may also trigger an immune response in mosquitoes is currently unknown. In a companion paper investigating *P. relictum* gene expression in this experiment (Sekar et al. 2021), we found a high P25 expression both in bird blood (immediately before the mosquito blood meal) and within the mosquito at 30 mpi, but very low expression at later timepoints (see Sekar et al. 2021, gene PRELSG_0614600). Second, the mosquito may (also) be reacting to the plasma of infected hosts. Plasma from *Plasmodium*-infected hosts contain several components of the host’s immune response (e.g. IgG, IgE and IgM antibodies, and TNF-α and related inflammatory cytokines) as well as extracellular vesicles shuttling communication signals between parasites during an infection (Clark 2007; Opadokun and Rohrbach 2021). When injected to a new vertebrate host, plasma from *Plasmodium*-infected individuals can exert powerful immune responses (Couper et al. 2010), though whether it can trigger a similar response in mosquitoes remains to be tested. Several other immune components (e.g. Cecropin B) were, in contrast, significantly down-regulated at 30 min post blood meal, a phenomenon that is suggestive of an immunomodulatory effect of the early stages of the parasite’s infection in the mosquito. *P. falciparum* Pfs47, which is expressed in gametes and early ookinetes, actively supresses a key immune signalling cascade that ultimately leads to the TEP1-mediated lysis of the parasite (Molina-Cruz et al. 2013; Ramphul et al. 2015; Molina-Cruz et al. 2020). In *P. relictum*, a Pfs47 orthologue was found to have high expression levels in bird blood as well as in the mosquito 30min post-infection, with nearly non existing expression at later time points (see Sekar et al. 2021, gene PRELSG_1251100).

Towards the end of the infection (22 dpi) the differences in immune gene expression between infected and uninfected mosquitoes are drastically reduced. Time transition analyses revealed that the lack of differences in both Toll and Imd pathways towards the end of the mosquito lifespan was due to an increase in immune gene expression in control mosquitoes rather than to a decrease in infected ones. This increase in immune investment towards the end of the mosquito life is paradoxical, as the general expectation is for a decline in immune function with age (immune senescence) as a result of physiological wear and tear, or of an adaptive reallocation of immune resources towards other traits such as reproduction or longevity (Shanley et al. 2009). Our results, disagree with previous results obtained in the *Cx pipiens/P relictum* system, that showed a decrease in PO activity (Cornet and Sorci 2010) and haematocyte counts (Pigeault et al. 2015) with age, as well as a higher susceptibility to a *P. relictum* infection in old mosquitoes (Pigeault et al. 2015). An increase in immune gene transcripts with age has however also been observed in *Drosophila melanogaster* (Zerofsky et al. 2005), which has been tentatively interpreted as the result of an age-related failure of one of the myriad mechanisms responsible for regulating immune response (Zerofsky et al. 2005), highlighting the limitations of extrapolating immune function from immune transcript quantification.

In conclusion, the present study shows that Toll and Imd immune pathways appear to be important over the infection through the up regulation of several receptors, translation factors and effectors previously identified in *Anopheles* mosquitoes infected with human and rodent malaria. By comparing infected and uninfected mosquitoes at different time-points after the bloodmeal we also revealed a dynamic pattern of gene expression that raises interesting questions about the onset of *Plasmodium* immunogenicity and mosquito immunosenescence, showcasing the added value of temporal transcriptomic studies of *Plasmodium*-mosquito interactions.

## MATERIAL AND METHODS

### Experimental design

The aim of this experiment is to understand how avian malaria infections influence expression patterns of immune genes in mosquitoes at four different key stages of the *P. relictum* development within the mosquito (Sekar et al. 2021): 30 minutes after the blood meal ingestion (gametocyte activation and formation of gametes), 8 days post infection (peak of oocyst production), 12 days post infection (peak sporozoite production) and 22 days post infection (ending stages of the infection). For this purpose, 3 canaries were infected with a standard dose of *P. relictum* (lineage pSGS1) following previously-publish protocols (Pigeault, Vézilier, et al. 2015), 3 other canaries were used as uninfected controls.

Ten days later, at the peak of the acute infection stage in blood (Valkiūnas 2005), the 3 infected and 3 control birds were placed individually in an experimental cage with 150 seven-day old female mosquitoes, which had been reared using standard laboratory protocols (Vézilier et al. 2010). Cages were visited 30 minutes later to take out the bird and any mosquitoes that were not fully gorged. At this point, 10 fully-gorged resting mosquitoes were haphazardly sampled from each of the cages (‘30mpi’ sample), homogenised with 500 μl of TRIzol LS^™^, and frozen at -80°C for subsequent RNA extraction (1 pool of 10 mosquitoes per cage). The rest of the mosquitoes were left in their cages with a source of sugar solution (10%) at our standard insectary conditions (25-27°C, 70% RH). On day 8 after the blood meal, 10 mosquitoes were randomly taken from each of the three cages (‘8 dpi’ sample), homogenised with 500 μl of RNAlater^®^ and frozen at -80°C. The procedure was repeated on days (‘12 dpi’ sample) and 22 (‘22 dpi’ sample). TRIzol was used for the blood engorged mosquitoes because bird blood, with its nucleated red blood cells, clogs the filters of the RNA extraction spin columns if they aren’t first treated in a TRIzol step (see below).

### RNA extraction

All analyses were carried out using pools of 10 mosquitoes (1 pool per time point per bird). For the 30mpi samples, the total volume of the buffer was adjusted to 750 μl; the sample, containing mosquitoes and the buffer, was subsequently homogenised using a TissueLyser (Qiagen, Hilden, Germany) equipped with a 5 mm stainless steel bead. The TissueLyser was run for two cycles of three minutes at 30 Hz. Phase separation was done according to the TRIzol LS^™^ manufacturer’s protocol; the resulting aqueous phase was mixed with one volume of 70% ethanol and placed in a RNeasy Mini spin column. RNA from the 8dpi, the 12dpi and 22dpi samples was extracted by first transferring the mosquitoes to a new tube together with 600 μl of buffer RLT and a 5 mm stainless steel bead and then homogenized using a TissueLyser. The TissueLyser was run for two cycles of three minutes at 30 Hz. Afterwards RNA was extracted using RNeasy Mini spin columns following the manufacturer’s protocol.

The concentration of all RNA samples was measured on a Nanodrop 2000/2000c (Thermo Fisher Scientific, Wilmington, DE, USA). mRNA from each time point was sequenced using Illumina HiSeq platform at an average of 85 M reads per library (Novogene, Hong Kong). We obtained paired-end reads of 150bp length.

### Data processing

Sequence data quality testing was performed using FASTQC (v 0.11.8) (Simon Andrews 2020). Low-quality reads were filtered or trimmed with Trimmomatic (v 0.27) (Bolger et al. 2014). The resulting files were aligned with STAR (v 2.7.9a) (Dobin et al. 2013) by using the *C. quinquefascistus* genome as reference (Holt et al. 2002). Finally, read count per gene was performed using FeatureCounts (v 2.0.1.1) (Liao et al. 2014).

### Statistical analyses

All the statistical analyses were carried out with the free statistical software R (R Core Team (2020) and the free integrated development environment RStudio (Rstudio 2020). The package DESeq2 (Version 1.16.1) (Love et al. 2014) was used to estimate the variance-mean dependence in count data from high-throughput sequencing assays and to test for differential expression based on a model using the negative binomial distribution. When testing for significant difference in expression, and in order to avoid problems arising from sequencing depth, gene length or RNA composition, the count data was first normalized in DESeq2 (Bushel et al. 2020). Principal component analyses (PCA) were built with a DESeq2 function for plotting PCA plots, that uses ggplot2 (Wilkinson 2011) under the hood.

For the purpose of investigating how different immune-pathways are activated in the mosquito during the infection, a list of genes was generated for each pathway through Vector Base bioinformatic resource (Amos et al. 2021) and employed for DeSeq2 analyses after the normalization process and statistical analyses that were done using the full gene-set. At each time point, the expression levels of the immune-genes were compared between infected and control mosquitoes (see Figure 2).

To further investigate the drastic decrease in differential expression of all gene pathways towards the end of the infection (day 22) we analysed the differences in gene expression occurring across the different time transitions (30mpi – 8dpi, 8dpi – 12dpi and 12dpi – 22dpi) in infected and control mosquitoes. This analysis did not distinguish between immune pathways.

### Data accessibility

Sequences have been uploaded to the Sequence Read Archive (SRA) at NCBI under accession number: PRJNA848963.

## ACKNOWLEDGMENTS AND FUNDING INFORMATION

Funding was provided by the Junta de Extremadura (PO17024, Post-Doc grant) to LGL; FEDER / Consejería de Economía, Ciencia y Agenda Digital, reference IB20089 to LGL. Swedish Research Council (grant 2016-03419 and 2021-03663) and Nilsson-Ehle foundation to OH. and the Agence Nationale de Recherche: ANR-16-CE35-0001-01 (‘EVODRUG’) to AR.

## LEGENDS FOR SUPPLEMENTARY MATERIAL

**Figure S1.**
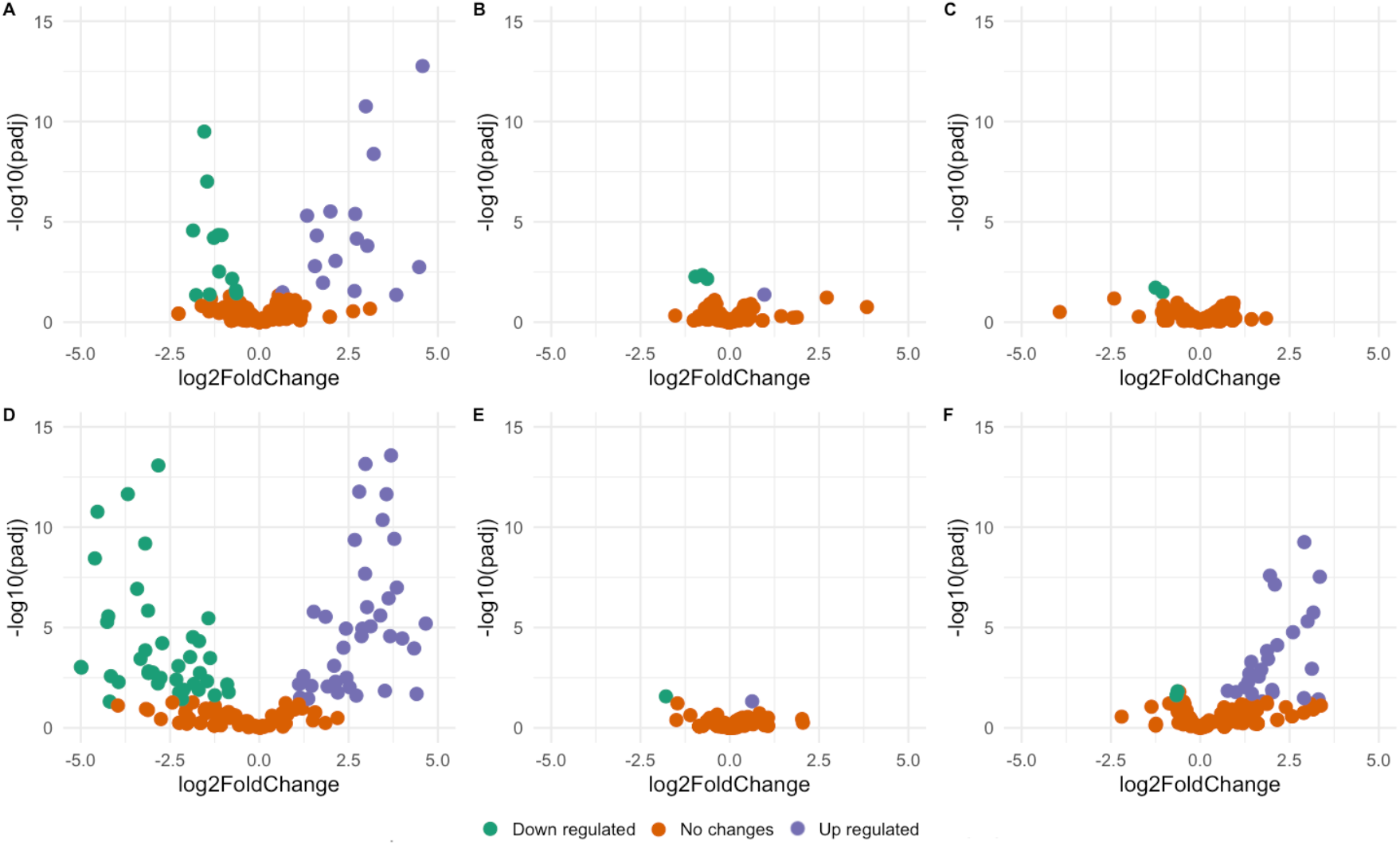
Volcano plots of differences in gene expression occurring across the different time transitions (30mpi – 8dpi, 8dpi – 12dpi and 12dpi – 22dpi) in infected and control mosquitoes: 30 mpi -8 days in infected mosquitos (A) and control mosquitos (D); 8 dpi – 12 dpi in infected (B) and control mosquitos (E); 12 dpi – 22 dpi in infected (C) and control mosquitos (F). Significant up-regulated (purple) and significant downregulated (green) dots were calculated using an adjusted p threshold of p = 0.05. Orange dots indicate no significant difference.

**Table S1**. Statistic results when comparing RNA expression of infected versus control mosquitos for the three immune pathways (Toll, Ims and JAK/STAT) at the different sampling time: 30 min (A), 8 days (B), 12 days (C) and 22 days (D).

**Table S2**. Functions and names for all genes included in the statistics. Genes were classified in the three immune pathways (Toll, Imd and JAK/STAT) by following a list of genes generated for each pathway through Vector Base bioinformatic resource (Amos et al. 2021).

## REFERENCES

Amos B, Aurrecoechea C, Barba M, Barreto A, Basenko EY, Ba W, Belnap R, Blevins AS, Ulrike B, Brestelli J, et al. 2021. VEuPathDB : the eukaryotic pathogen, vector and host bioinformatics resource center. :1–14.

Bartholomay LC, Waterhouse RM, Mayhew GF, Campbell CL, Michel K, Zou Z, Ramirez JL, Das S, Alvarez K, Arensburger P, et al. 2010. Pathogenomics of Culex quinquefasciatus and meta-analysis of infection responses to diverse pathogens. Science (1979).

Boëte C. 2005. Malaria parasites in mosquitoes: laboratory models, evolutionary temptation and the real world. Trends in Parasitology 21:445–447.

Bolger AM, Lohse M, Usadel B. 2014. Trimmomatic: A flexible trimmer for Illumina sequence data. Bioinformatics 30:2114–2120.

Bueno MG, Lopez RPG, Menezes RMTD, Costa-Nascimento MDJ, Lima GFMDC, Araújo RADS, Guida FJ v, Kirchgatter K. 2010. Identification of Plasmodium relictum causing mortality in penguins (Spheniscus magellanicus) from São Paulo Zoo, Brazil. Veterinary Parasitology 173:123–127.

Bushel PR, Ferguson SS, Ramaiahgari SC, Paules RS, Auerbach SS. 2020. Comparison of Normalization Methods for Analysis of TempO-Seq Targeted RNA Sequencing Data. Frontiers in Genetics 11.

Cao X, Gulati M, Jiang H. 2017. Serine protease-related proteins in the malaria mosquito, Anopheles gambiae. Insect Biochemistry and Molecular Biology 88:48–62.

del Carmen Rodriguez M, Gerold P, Dessens J, Kurtenbach K, Schwartz RT, Sinden RE, Margos G. 2000. Characterisation and expression of pbs25, a sexual and sporogonic stage specific protein of Plasmodium berghei. Mol Biochem Parasitol 110:147–159.

Carr AL, Rinker DC, Dong Y, Dimopoulos G, Zwiebel LJ. 2021. Transcriptome profiles of Anopheles gambiae harboring natural low-level Plasmodium infection reveal adaptive advantages for the mosquito. Scientific Reports 11:1–25.

Cellier-Holzem E, Esparza-Salas R, Garnier S, Sorci G. 2010. Effect of repeated exposure to Plasmodium relictum (lineage SGS1) on infection dynamics in domestic canaries. International Journal for Parasitology 40:1447–1453.

Challenger JD, Olivera Mesa D, Da DF, Yerbanga RS, Lefèvre T, Cohuet A, Churcher TS. 2021. Predicting the public health impact of a malaria transmission-blocking vaccine. Nature Communications.

Cirimotich CM, Dong Y, Garver LS, Sim S, Dimopoulos G. 2010. Mosquito immune defenses against Plasmodium infection. Developmental and Comparative Immunology 34:387–395.

Clark IA. 2007. How TNF was recognized as a key mechanism of disease. Cytokine Growth Factor Rev 18:335–343.

Clayton AM, Dong Y, Dimopoulos G. 2014. The anopheles innate immune system in the defense against malaria infection. Journal of Innate Immunity 6:169–181.

Cornet S, Sorci G. 2010. Parasite virulence when the infection reduces the host immune response. Proceedings. Biological sciences / The Royal Society 277:1929–1935.

Couper KN, Barnes T, Hafalla JCR, Combes V, Ryffel B, Secher T, Grau GE, Riley EM, de Souza JB. 2010. Parasite-derived plasma microparticles contribute significantly to malaria infection-induced inflammation through potent macrophage stimulation. PLOS Pathogens 6:e1000744.

Dobin A, Davis CA, Schlesinger F, Drenkow J, Zaleski C, Jha S, Batut P, Chaisson M, Gingeras TR. 2013. STAR: Ultrafast universal RNA-seq aligner. Bioinformatics 29:15–21.

Dong Y, Aguilar R, Xi Z, Warr E, Mongin E, Dimopoulos G. 2006. Anopheles gambiae immune responses to human and rodent Plasmodium parasite species. PLoS Pathogens 2(6)e52.

Eckhoff PA. 2011. A malaria transmission-directed model of mosquito life cycle and ecology. Malaria Journal 10(1):1–17.

Garver LS, Dong Y, Dimopoulos G. 2009. Caspar controls resistance to Plasmodium falciparum in diverse anopheline species. PLoS Pathogens 5(3):e10010335.

Girard YA, Mayhew GF, Fuchs JF, Li H, Schneider BS, McGee CE, Rocheleau TA, Helmy H, Christensen BM, Higgs S, et al. 2010. Transcriptome changes in Culex quinquefasciatus (Diptera: Culicidae) salivary glands during west nile virus infection. Journal of Medical Entomology 47(3):421–435.

Hajkazemian M, Bossé C, Mozūraitis R, Emami SN. 2021. Battleground midgut: The cost to the mosquito for hosting the malaria parasite. Biology of the Cell 113(2):79–94.

Health W, Who O. 2021. WHO Guidelines for malaria. Geneva: World Health Organization. Letters in Applied Microbiology.

Hellgren O, Atkinson CT, Bensch S, Albayrak T, Dimitrov D, Ewen JG, Kim KS, Lima MR, Martin L, Palinauskas V, et al. 2015. Global phylogeography of the avian malaria pathogen Plasmodium relictum based on MSP1 allelic diversity. Ecography 38:842–850.

Holt RA, Mani Subramanian G, Halpern A, Sutton GG, Charlab R, Nusskern DR, Wincker P, Clark AG, Ribeiro JMC, Wides R, et al. 2002. The genome sequence of the malaria mosquito Anopheles gambiae. Science (1979) 298:129–149.

Howick VM, Russell AJC, Andrews T, Heaton H, Reid AJ, Natarajan K, Butungi H, Metcalf T, Verzier LH, Rayner JC, et al. 2019. The malaria cell atlas: Single parasite transcriptomes across the complete Plasmodium life cycle. Science (1979).

Kanehisa M, Sato Y, Kawashima M. 2021. KEGG mapping tools for uncovering hidden features in biological data. Protein Science.

Kar NP, Kumar A, Singh OP, Carlton JM, Nanda N. 2014. A review of malaria transmission dynamics in forest ecosystems. Parasites and Vectors 7(1):1–12.

Kazlauskiene R, Bernotiene R, Palinauskas V, Iezhova TA, Valkiunas G. 2013. Plasmodium relictum (lineages pSGS1 and pGRW11): Complete synchronous sporogony in mosquitoes Culex pipiens pipiens. Experimental Parasitology 133:454–461.

Keleta Y, Ramelow J, Cui L, Li J. 2021. Molecular interactions between parasite and mosquito during midgut invasion as targets to block malaria transmission. npj Vaccines 6:1–9.

Kokoza V, Ahmed A, Shin SW, Okafor N, Zou Z, Raikhel AS. 2010. Blocking of Plasmodium transmission by cooperative action of Cecropin A and Defensin A in transgenic Aedes aegypti mosquitoes. Proc Natl Acad Sci U S A 107:8111– 8116.

Kumar S, Christophides GK, Cantera R, Charles B, Han YS, Meister S, Dimopoulos G, Kafatos FC, Barillas-Mury C. 2003. The role of reactive oxygen species on Plasmodium melanotic encapsulation in Anopheles gambiae. Proc Natl Acad Sci U S A 100:14139–14144.

Liao Y, Smyth GK, Shi W. 2014. FeatureCounts: An efficient general purpose program for assigning sequence reads to genomic features. Bioinformatics 30:923–930.

Lorenz C, Alves JMP, Foster PG, Suesdek L, Sallum MAM. 2021. Phylogeny and temporal diversification of mosquitoes (Diptera: Culicidae) with an emphasis on the Neotropical fauna. Systematic Entomology 46:798–811.

Love M, Anders S, Huber M. 2014. Differential gene expression analysis based on the negative binomial distribution. Genome Biol 15, 550.

Luckhart S, Vodovotz Y, Ciu L, Rosenberg R. 1998. The mosquito Anopheles stephensi limits malaria parasite development with inducible synthesis of nitric oxide. Proc Natl Acad Sci U S A 95:5700–5705.

Molina-Cruz A, Canepa GE, Alves E Silva TL, Williams AE, Nagyal S, Yenkoidiok-Douti L, Nagata BM, Calvo E, Andersen J, Boulanger MJ, et al. 2020. Plasmodium falciparum evades immunity of anopheline mosquitoes by interacting with a Pfs47 midgut receptor. Proc Natl Acad Sci U S A 117:2597– 2605.

Molina-Cruz A, Garver LS, Alabaster A, Bangiolo L, Haile A, Winikor J, Ortega C, van Schaijk BCL, Sauerwein RW, Taylor-Salmon E, et al. 2013. The human malaria parasite Pfs47 gene mediates evasion of the mosquito immune system. Science (1979) 340:984–987.

Moyes CL, Athinya DK, Seethaler T, Battle KE, Sinka M, Hadi MP, Hemingway J, Coleman M, Hancock PA. 2021. Evaluating insecticide resistance across African districts to aid malaria control decisions. PNAS 36:117.

Onyango SA, Ochwedo KO, Machani MG, Omondi CJ, Debrah I, Ogolla SO, Lee MC, Zhou G, Kokwaro E, Kazura JW, et al. 2021. Genetic diversity and population structure of the human malaria parasite Plasmodium falciparum surface protein Pfs47 in isolates from the lowlands in Western Kenya. PLOS ONE 16:e0260434.

Opadokun T, Rohrbach P. 2021. Extracellular vesicles in malaria: an agglomeration of two decades of research. Malaria Journal 20:1–16.

Peterson TML, Gow AJ, Luckhart S. 2007. Nitric oxide metabolites induced in Anopheles stephensi control malaria parasite infection. Free Radic Biol Med 42:132.

Pigeault R, Nicot A, Gandon S, Rivero A. 2015. Mosquito age and avian malaria infection. Malaria Journal 14:1–11.

Pigeault R, Vézilier J, Cornet S, Zélé F, Nicot A, Perret P, Gandon S, Rivero A. 2015. Avian malaria: A new lease of life for an old experimental model to study the evolutionary ecology of Plasmodium. Philosophical Transactions of the Royal Society B: Biological Sciences 370.

R Core Team (2020). 2020. R: A language and environment for statistical computing. R: A language and environment for statistical computing. R Foundation for Statistical Computing, Vienna, Austria.

Ramphul UN, Garver LS, Molina-Cruz A, Canepa GE, Barillas-Mury C. 2015. Plasmodium falciparum evades mosquito immunity by disrupting JNK-mediated apoptosis of invaded midgut cells. Proc Natl Acad Sci U S A 112:1273–1280.

Raulf MK, Johannssen T, Matthiesen S, Neumann K, Hachenberg S, Mayer-Lambertz S, Steinbeis F, Hegermann J, Seeberger PH, Baumgärtner W, et al. 2019. The C-type Lectin Receptor CLEC12A Recognizes Plasmodial hemozoin and contributes to cerebral malaria development. Cell Reports 28:30-38.e5.

Reynolds RA, Kwon H, Smith RC. 2020. 20-Hydroxyecdysone Primes Innate Immune Responses That Limit Bacterial and Malarial Parasite Survival in Anopheles gambiae. Msphere 5(2) e00983–19.

Rivero A, Gandon S. 2018. Evolutionary Ecology of Avian Malaria: Past to Present. Trends in Parasitology 34:712–726.

Rstudio T. 2020. RStudio: Integrated Development for R. Rstudio Team, PBC, Boston, MA URL http://www.rstudio.com/.

Ruiz JL, Yerbanga RS, Lefèvre T, Ouedraogo JB, Corces VG, Gómez-Díaz E. 2019. Chromatin changes in Anopheles gambiae induced by Plasmodium falciparum infection. Epigenetics and Chromatin 12:1–18.

Ryan SJ, Lippi CA, Zermoglio F. 2020. Shifting transmission risk for malaria in Africa with climate change: A framework for planning and intervention. Malaria Journal 19(1):1–14.

Saxena AK, Wu Y, Garboczi DN. 2007. Plasmodium P25 and P28 surface proteins: Potential transmission-blocking vaccines. Eukaryotic Cell 6:1260–1265.

Sekar V, Rivero A, Pigeault R, Gandon S, Drews A, Ahren D, Hellgren O. 2021. Gene regulation of the avian malaria parasite Plasmodium relictum, during the different stages within the mosquito vector. Genomics 113(4):2327–2337.

Shanley DP, Aw D, Manley NR, Palmer DB. 2009. An evolutionary perspective on the mechanisms of immunosenescence. Trends in Immunology 30:374–381.

Shin D, Civana A, Acevedo C, Smartt CT. 2014. Transcriptomics of differential vector competence: West Nile virus infection in two populations of Culex pipiens quinquefasciatus linked to ovary development. BMC Genomics.

da Silva AF, Machado LC, de Paula MB, da Silva Pessoa Vieira CJ, de Morais Bronzoni RV, de Melo Santos MAV, Wallau GL. 2020. Culicidae evolutionary history focusing on the Culicinae subfamily based on mitochondrial phylogenomics. Scientific Reports 2020 10:1–14.

Simões ML, Dong Y, Mlambo G, Dimopoulos G. 2022. C-type lectin 4 regulates broad-spectrum melanization-based refractoriness to malaria parasites. PLOS Biology 20:e3001515.

Simon Andrews. 2020. Babraham Bioinformatics - FastQC A Quality Control tool for High Throughput Sequence Data. Soil 5:47–81.

Singh M, Suryanshu, Kanika, Singh G, Dubey A, Chaitanya RK. 2021. Plasmodium’s journey through the Anopheles mosquito: A comprehensive review. Biochimie 181:176–190.

Tahar R, Boudin C, Thiery I, Bourgouin C. 2002. Immune response of Anopheles gambiae to the early sporogonic stages of the human malaria parasite Plasmodium falciparum. The EMBO Journal 21:6673–6680.

Tikhe C V., Dimopoulos G. 2021. Mosquito antiviral immune pathways. Developmental and Comparative Immunology 116, 103964.

Valkiunas G. 2005. Avian malaria parasites and other Haemosporidia. Boca Raton. CRC press.

Valkiunas G, Iezhova TA. 2018. Keys to the avian malaria parasites. Malaria Journal 17(1):1–24.

Valkiunas G, Ilgunas M, Bukauskaite D, Fragner K, Weissenböck H, Atkinson CT, Iezhova TA. 2018. Characterization of Plasmodium relictum, a cosmopolitan agent of avian malaria. Malaria Journal 17(1):1–21.

Vargas V, Cime-Castillo J, Lanz-Mendoza H. 2020. Immune priming with inactive dengue virus during the larval stage of Aedes aegypti protects against the infection in adult mosquitoes. Scientific Reports 2020 10:1

Vézilier J, Nicot A, Gandon S, Rivero A. 2010. Insecticide resistance and malaria transmission: Infection rate and oocyst burden in Culex pipiens mosquitoes infected with Plasmodium relictum. Malaria Journal 9.

Wadi I, Anvikar AR, Nath M, Pillai CR, Sinha A, Valecha N. 2018. Critical examination of approaches exploited to assess the effectiveness of transmission-blocking drugs for malaria. Future Medicinal Chemistry 10(22):2619–2639.

Wilkinson L. 2011. ggplot2: Elegant Graphics for Data Analysis by WICKHAM, H. Biometrics 67:678–679.

Zerofsky M, Harel E, Silverman N, Tatar M. 2005. Aging of the innate immune response in Drosophila melanogaster. Aging Cell 4:103–108.

Zhang X, An C, Sprigg KJ, Michel K. 2016. CLIPB8 is part of the prophenoloxidase activation system in Anopheles gambiae mosquitoes. Insect Biochemistry and Molecular Biology 71:106–115.

